# Tricotyledonous Seedling in Jute (*Corchorus capsularis*): First Observation During Tissue Culture Selection

**DOI:** 10.64898/2025.12.28.696764

**Authors:** Subhadarshini Parida, Shuvobrata Majumder

## Abstract

White jute (*Corchorus capsularis*) is a dicotyledonous, annual, bast fiber-producing crop. Under controlled culture room conditions (27 ± 2°C, 16 h light/8 h dark photoperiod) and in a 15 mg/L hygromycin-B supplemented nutrient medium, one out of 50 tested seeds (2%) exhibited a rare tricotyledonous seedling phenotype. This is the first documented occurrence of tricotyledon in white jute during tissue culture selection of the cultivar JRC321 (Sonali). Given the potential advantages of polycotyledon in other plant species—such as improved vigor, and biomass accumulation—this finding opens new avenues for investigating the genetic and physiological mechanisms underlying tricotyledon development in jute. Future studies could explore its implications for breeding programs aimed at enhancing fiber yield and production.

## 1. Introduction

Jute is a major bast fiber crop, cultivated extensively in India, Bangladesh, and China, to meet the global demand for its long, shiny, and biodegradable fibers. Jute belongs to the Malvaceae family, which comprises approximately 4,225 species. Interestingly, among these, only seven species (*Abutilon theophrasti, Alcea rosea, Gossypium hirsutum, Hibiscus* sp., *Tilia cordata subsp. cordata, Tilia europaea*, and *Corchorus capsularis*) have been reported to exhibit polycotyledons—a rare developmental anomaly where seedlings develop more than two cotyledons (**Fu et al., 2024**). Only two jute species are cultivated commercially: white jute (*Corchorus capsularis*) and tossa jute (*Corchorus olitorius*). A prior report on polycotyledonous white jute (*C. capsularis*) was published 71 years ago (**Datta, 1954**). However, we were unable to obtain a copy of this report, and no images of tricotyledonous seedlings in jute are available in digital media or online.

Cotyledons play a crucial role in seedling development, establishment, and survival, serving as an initial nutrient source and functioning as the first photosynthetic organ (**Chandle, 2008**). A recent study by **Fu et al. (2024)** reported that among 1,71,983 dicotyledonous angiosperm species, only 342 species exhibited polycotyledony. In contrast, 160 out of 618 gymnosperm species displayed this trait, indicating that polycotyledony is more frequent in gymnosperms than in angiosperms. The highest occurrence has been observed in the Pinaceae family (122 out of 250 species), followed by the Proteaceae family (25 out of 1,660 species). The number of cotyledons varies between 3 to 24 in gymnosperms, whereas in angiosperms, it typically ranges from 3 to 4. The exact causes of polycotyledony remain unknown, but it has been observed to occur spontaneously in natural populations at frequencies of up to 6% (**Ran et al., 2024; Fu et al., 2024**).

During our research on transgenic jute development, we observed an unusual case of polycotyledony in the white jute cultivar JRC321. A seedling exhibited an additional cotyledon with degraded chlorophyll (yellowing) under hygromycin-B selection. This represents the first documented case of polycotyledony in jute. The occurrence of this rare trait under selection pressure raises important questions about its underlying genetic and environmental influences. Further studies are required to understand the molecular mechanisms governing cotyledon development in jute.

## 2. Materials and Methods

### 2.1. Preparation of Germination Medium

For seed germination and transgenic jute seed selection, a culture medium was prepared using Murashige and Skoog (MS) basal salts and vitamins (HiMedia Lab. Pvt. Ltd., Mumbai, India), supplemented with 15% sucrose and 15 mg/L Hygromycin-B as a selective agent. The medium was solidified with 0.8% agar and adjusted to pH 5.8 before autoclaving (**Majumder et al., 2018a**).

### 2.2. Seed Surface Sterilization and Germination

White jute seeds (cultivar JRC321) were surface-sterilized by sequentially treating them with 0.1% Bavistin (carbendazim 50%) fungicide for 10 min, followed by 70% ethanol for 4 min, and then gently agitating them in a sodium hypochlorite solution for 15 min (**Majumder et al., 2024a**). After sterilization, the seeds were thoroughly rinsed five times with sterile distilled water to remove any residual chemicals. The sterilized seeds were then placed on sterile filter paper to remove excess moisture before being transferred to the prepared germination medium. Seed germination was initiated in a 37°C incubator in complete darkness for 48 h, after which the seedlings were moved to a growth chamber maintained at 27 ± 2°C under a 16-h light / 8-h dark photoperiod for further development.

### 2.3. Calculation of Germination and Polycotyly Percentages

Seed germination percentage was determined seven days after sowing using the formula: (Number of germinated seedlings ÷ Total number of seeds in the plates) ⍰ 100. Similarly, the polycotyledony percentage was calculated based on the proportion of seedlings exhibiting an additional cotyledon, using the formula: (Number of seedlings with three cotyledons ÷ Total number of germinated seedlings) ⍰ 100.

## 3. Results

The majority (96%) of seeds germinated successfully, with seedlings exhibiting a mix of healthy and affected phenotypes (**Figure 1A**). Seedlings exposed to 15 mg/L hygromycin-B displayed characteristic stress symptoms, including stem and root browning, chlorophyll degradation in cotyledons (resulting in yellowing), stunted growth, reduced root development, and necrosis.

**Figure 1.**
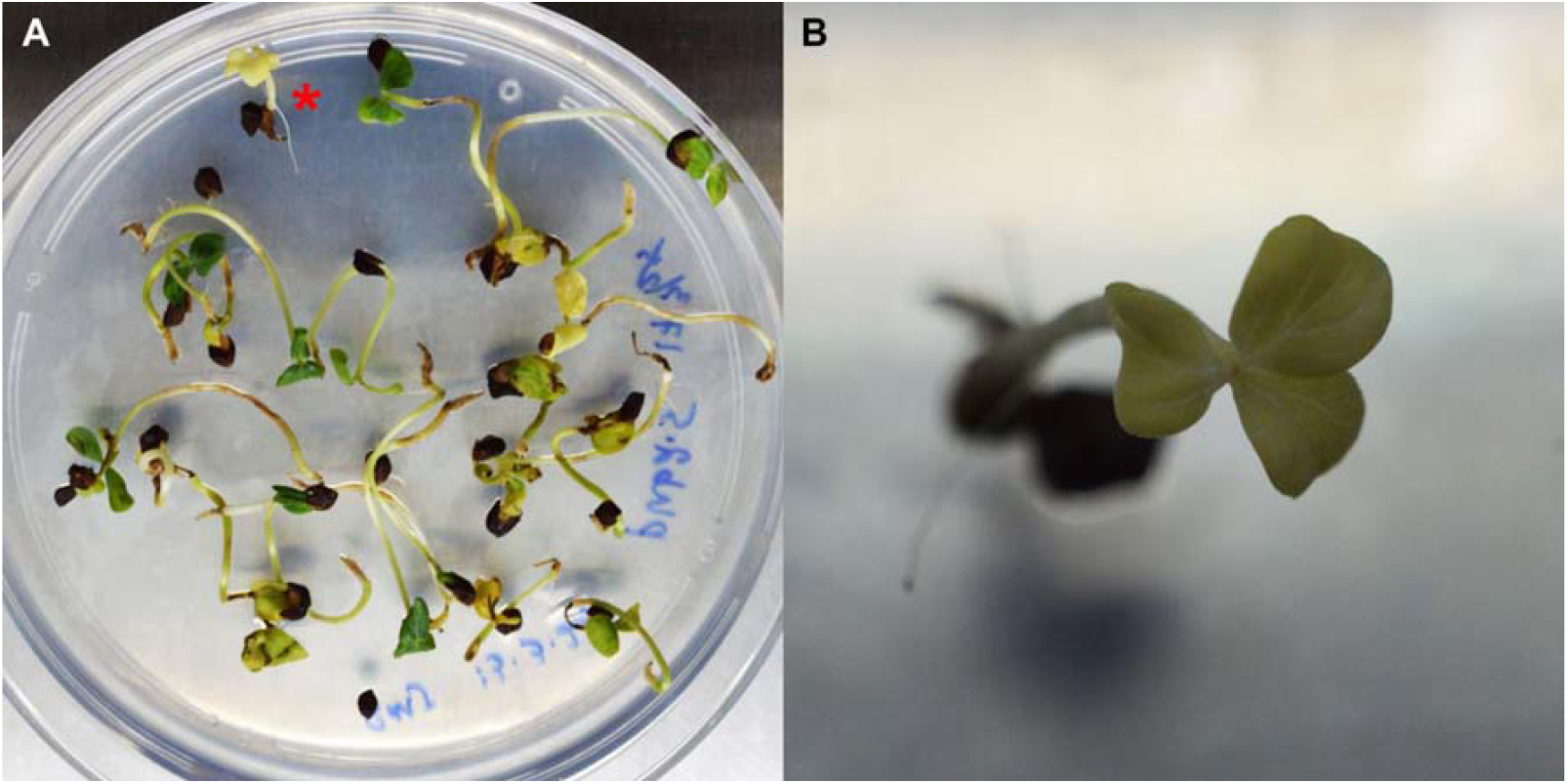
Seedling phenotypes of white jute cultivar JRC321 grown on hygromycin-B (15 mg/L) supplemented medium. (A) Seedlings after 7 days under hygromycin-B selection stress, showing a mix of healthy and affected phenotypes. The seedling exhibiting tricotyledon is marked with a red asterisk. (B) Close-up view of the tricotyledonous seedling.

Among all germinated seedlings, only one exhibited an unusual tricotyledonous phenotype, while the rest developed the typical dicotyledonous structure (**Figure 1B**). The observed frequency of tricotyledon development was 2%.

## 4. Discussion

Polycotyledony is a rare but naturally occurring phenomenon observed in multiple plant species (**Holtorp, 1944**). A recently published review by **Fu et al. (2024)** compiled a list of 160 species from six gymnosperm families and 342 species from 80 angiosperm families exhibiting polycotyly, including a previous report on a tricotyledonous white jute seedling. The earlier report by **Datta (1954)** on jute tricotyledony is over seven decades old and is not currently available in any accessible media. Over the past decades, concepts and understanding of polycotyledony have evolved, yet some aspects remain unclear, with scientific explanations still pending. In this section, we discuss these aspects while highlighting our findings in jute crops.

How do polycots develop? Previous reports suggest that polycots can occur spontaneously (**Ran et al., 2024**), be induced by hybridization (**Stubbe, 1963**) or mutagenesis (**Haccius, 1955; Zhang and Wei, 1990**), and be influenced by environmental factors such as highly polluted sites (**Weiersbye and Witkowski, 2003**), common gardens (**DeVries, 1909; 1910**), different climates (**Harrison, 1964; Palmer, 1968**), and seasonal variations (**Sargent, 1903**). Additionally, polycotyledony has been observed in response to tissue culture media (**Homes and Guillaume, 1967**) and even the positional arrangement of fruits on a plant, such as in tomatoes (**Haskell, 1954**). Among the possible influencing factors, our study on jute suggests that tricotyledon development may have occurred either spontaneously or as a response to tissue culture media supplemented with hygromycin-B. Other potential factors can be ruled out, as the jute plants were grown under controlled conditions (27 ± 2°C under a 16-h light / 8-h dark photoperiod).

Since 2014, we have used hygromycin-B supplemented media for the selection of transgenic white jute (*C. capsularis*) cultivar JRC321, starting with our first report on *Agrobacterium*-mediated transformation standardization using shoot-tip explants (**Saha et al., 2014**) and more recently with imbibed seed explants (**Majumder et al., 2024a**). Over the past decade, hygromycin-B has been consistently used for developing multiple transgenic traits in white jute, including insect resistance (**Majumder et al., 2025**) and pyramided multi-trait resistance to insects, fungi, and herbicides (**Majumder et al., 2024b**). However, we have never previously observed tricotyledon development. Similarly, no instances of tricotyledony were recorded under other selection pressures, such as kanamycin (**Majumder et al., 2020**) or the herbicide compound glufosinate ammonium (**Majumder et al., 2018b**). These selection agents (hygromycin-B, kanamycin, and glufosinate ammonium) are used to identify transgenic plants while eliminating non-transgenic ones, which exhibit phenotypic traits such as stem and root browning, rapid degradation of cotyledon chlorophyll, stunted growth, lack of lateral root formation, and necrosis (**Majumder et al., 2018a**). Non-transgenic seeds typically fail to germinate on selection media. While these phenotypic responses can be seen in **Figure 1**, tricotyledony development has never been observed during tissue culture until now. This is the first recorded instance of tricotyledony in JRC321, suggesting that it likely occurred spontaneously at a very low frequency (2%) rather than being induced by hygromycin-B.

Tricotyledony has been reported to offer agricultural advantages over dicotyledony in various following plant species. For instance, in sugar beet (*Beta vulgaris*), it is linked to larger seed clusters; in castor oil plant (*Ricinus communis*), tricotyledonous individuals exhibit greater vigor and fertility compared to dicotyledonous ones (**Haskell et al., 1954; Litovchenko, 1940**). Similarly, in Mexican cypress (*Cupressus lusitanica*), tricotyledonous seedlings demonstrate improved growth and height, while in *Eucalyptus*, they tend to be more vigorous than their dicotyledonous counterparts (**Griffith, 1953; Venkatesh and Sharma, 1974**). Despite these observations, detailed studies exploring the specific agricultural or pharmaceutical implications of tricotyledony within individual species remain limited. In the case of jute, the potential benefits of an additional cotyledon have not yet been analyzed. However, if tricotyledony contributes to increased plant height and growth, it could represent a valuable trait for breeding programs aimed at enhancing jute productivity.

The genetic basis of tricotyledony remains an underexplored area of research. However, some studies in *Arabidopsis thaliana* have provided insights into cotyledon development, as reviewed by **Chandler (2008)**. Gene families such as *PIN* and *DRN/DRNL* have been identified in *Arabidopsis* as key regulators of cotyledon formation (**Friml et al., 2003; Chandler et al., 2007**). Current understanding suggests that cotyledon development is governed by multiple genes and pathways, with auxin playing a crucial role through the *STM* pathway. Additionally, a study on *Arabidopsis* mitochondrial iron transporter (*MIT*) genes revealed that mutations in *AtMIT1* and *AtMIT2* resulted in tricotyledony, which was considered an abnormal phenotype in the study (**Vargas et al., 2023**). However, no information is currently available on genes or pathways associated with tricotyledony in jute. This presents a significant opportunity for future research to uncover the genetic mechanisms underlying this rare trait in *C. capsularis*.

### CRediT authorship contribution statement

**Subhadarshini Parida**: Writing original draft, Investigation, Formal analysis, Data curation. **Shuvobrata Majumder**: Writing review & editing, Supervision, Visualization, Investigation, Funding acquisition.

## Declarations

### Ethical approval

Not applicable

### Clinical trial

Not applicable

## Acknowledgments

We express our gratitude to the late Dr. Ajay Parida, former Director, and Dr. Debasis Dash, Director of BRIC-Institute of Life Sciences, Bhubaneswar, Odisha, India, for their steadfast support and vital resources indispensable to the project’s success. We sincerely appreciate Mrs. Soma Roy from Bioingene.com for her insightful comments and suggestions as an internal reviewer.

## Data availability

The datasets generated during and/or analysed during the current study are available from the corresponding author on reasonable request.

## References

1. Fu Y-B. Polycotyly: How Little Do We Know? Plants. 2024;13:1054. 10.3390/plants13081054

2. Datta RM. Tricotyledony in Corchorus capsularis Linn. Science and Culture. 1954;20:240−241.

3. Chandler JW. Cotyledon organogenesis. J Exp Bot. 2008;59:2917–2931. 10.1093/jxb/ern167

4. Ran R, Li X, Sun H, Liu Y, Chen G, Zhao P. Tricotyledony in sand rice (Agriophyllum squarrosum). Genet Resour Crop Evol. 2024;71:529–537. 10.1007/s10722-023-01712-7

5. Majumder S, Sarkar C, Saha P, Gotyal BS, Satpathy S, Datta K, Datta SK. Bt jute expressing fused δ-endotoxin Cry1Ab/Ac for resistance to Lepidopteran pests. Front Plant Sci. 2018a;8:2188. 10.3389/fpls.2017.02188

6. Majumder S, Parida S, Dey N. Protocol for imbibed seed piercing for Agrobacterium-mediated transformation of jute. Star Protoc. 2024a;5:102767. 10.1016/j.xpro.2023.102767

7. Holtorp HE. Tricotyledony. Nature. 1944;153:13–14. 10.1038/153013a0

8. Stubbe H. Uber die Stabilisierung des sich variabel manifestierenden Merkmals Polycotylie von Antirrhinum malus. Die Kult. 1963;11:250–263. 10.1007/BF02136117

9. Haccius B. Experimentally induced twinning in plants. Nature. 1955;176: 355–356. 10.1038/176355a0

10. Zhang Y-J, Wei X-P. Induction of tricotyledon from embryos of Vitis vinifera L. Acta Botanica Boreali-Occidentalia Sinica. J Northwest Bot. 1990;10:228–231.

11. Weiersbye IM, Witkowski ET. Acid rock drainage (ARD) from gold tailings dams on the Witwatersrand Basin impacts on tree seed fate, inorganic content and seedling morphology. In Mine Water and the Environment, Proceedings of the 8th International Mine Water and the Environment Congress, Johannesburg, South Africa, 7–13 September 1998; Armstrong D, de Villiers AB, Kleinmann RLP, McCarthy TS, Norton PJ, Eds.; IMWA: Wendelstein, Germany. 2003;311–330.

12. DeVries H. (1909). The Mutation Theory: Experiments and Observations on the Origin of Species in the Vegetable Kingdom Vol. I: The Origin of Species by Mutation. Open Court Publishing Company: Chicago, IL, USA. 1909.

13. DeVries H. The Mutation Theory: Experiments and Observations on the Origin of Species in the Vegetable Kingdom Vol. II: The Origin of Varieties by Mutation; Open Court Publishing Company: Chicago, IL, USA.

14. Harrison BJ. Factors affecting the frequency of tricotyly in Antirrhinum majus. Nature. 1964;201:424. 10.1038/201424a0

15. Palmer TP. Inheritance of split cotyledons in swede turnips (Brassica napus). N Z J Bot. 1968;6:129–136. 10.1080/0028825X.1968.10429055

16. Sargent E. A theory of the origin of monocotyledons found on the structure of their seedlings. Ann Bot. 1903;17:1–92. 10.1093/oxfordjournals.aob.a088906

17. Homes JL, Guillaume M. Phénomènes D’organogenese dans des Cultures in vitro de Tissus de Carotte (Daucus carota L.). Bull Soc Roy Bot Belg. 1967;100:239–258.

18. Haskell G. Influence of position of truss on pleiocotyly in tomato. Nature. 1954;173:311.

19. Saha P, Datta K, Majumder S, Sarkar C, China SP, Sarkar SN,Sarkar D, Datta SK. Agrobacterium-mediated genetic transformation of commercial jute cultivar Corchorus capsularis cv. JRC 321 using shoot tip explants. Plant Cell Tissue Organ Cult. 2014;118:313–326. 10.1007/s11240-014-0484-6

20. Majumder S, Datta K, Datta SK. 25 Years of Pesticidal Cry1Ab/Ac Fusion Proteins in Crop Protection: Advances in Bt Crop Development, Target Pest Management, Safety, Environmental Impact, and Regulatory Frameworks. J Crop Health. 2025;77:55. 10.1007/s10343-025-01119-7

21. Majumder S, Datta K, Datta SK. Pyramided transgenic jute (Corchorus capsularis) with biotic stress resistance and herbicide tolerance. Ind Crops Prod. 2024b;208:117776. 10.1016/j.indcrop.2023.117776

22. Majumder S, Sarkar C, Datta K, Datta SK. Establishment of the ‘imbibed seed piercing’ method for Agrobacterium mediated transformation of jute and flax bast fibre crops via phloem-specific expression of the β-glucuronidase Gene. Ind Crops Prod. 2020;154:112620. 10.1016/j.indcrop.2020.112620

23. Majumder S, Datta K, Sarkar C, Saha SC, Datta SK. The Development of Macrophomina phaseolina (Fungus) Resistant and Glufosinate (Herbicide) Tolerant Transgenic Jute. Front Plant Sci. 2018b;9:920. 10.3389/fpls.2018.00920

24. Litovchenko AG. The role of tricotyledons in the socialist plant industry. C R Acad Sci. 1940;27:816–820.

25. Griffith AL. Hybridisation in the cypresses in East Africa. Emp For Rev. 1953;32:363.

26. Venkatesh CS, Sharma VK. Some unusual seedlings of Eucaplyptus: Their genetic significance and value in breeding. Silvae Genet. 1974;22:120–124.

27. Friml J, Vieten A, Sauer M, Weijers D, Schwarz H, Hamann T, Offringa R, Jürgens G. Efflux-dependent auxin gradients establish the apical-basal axis of Arabidopsis. Nature. 2003;426:147–153. 10.1038/nature02085

28. Chandler JW, Cole M, Flier A, Grewe B, Werr W. The AP2 transcription factors DORNRÖSCHEN and DORNRÖSCHEN LIKE redundantly control Arabidopsis embryo patterning via interaction with PHAVOLUTA. Development. 2007;134:1653–1662. 10.1242/dev.001016

29. Vargas J, Gómez I, Vidal EA, Lee CP, Millar AH, Jordana X, Roschzttardtz H. Growth Developmental Defects of Mitochondrial Iron Transporter 1 and 2 Mutants in Arabidopsis in Iron Sufficient Conditions. Plants. 2023;12(5):1176. 10.3390/plants12051176

